# Visualizing nucleation, condensation and propagation of β-tubulin folding in chaperonin TRiC

**DOI:** 10.1101/2024.10.13.618036

**Authors:** Yanyan Zhao, Michael F. Schmid, Wah Chiu

## Abstract

The folding nucleus (FN) initiates protein folding and enables an efficient folding pathway. Here we directly visualize the tubulin FN consisting of a nonnative, partially assembled Rossmann fold, in the closed chamber of human chaperonin TRiC. Chaperonin TRiC interacts with non-natively folded secondary structural elements, stabilizing the nucleus for transition into its first native domain. Through progressive folding, the unfolded sequence goes through drastic spatial arrangement in the TRiC chamber to sample the conformational space, mediated by the highly dynamic CCT tails. The observed presence of individual nonnative secondary structures first in the nonnative FN and then around the incrementally folded native domains supports the hypothesis that tubulin folding in TRiC is a hierarchical process of nucleation, condensation and propagation in cooperation with TRiC subunits.

## Introduction

A newly synthesized polypeptide needs to fold into its native conformation to function properly. Despite the seemingly astronomical number of possible conformations a protein can theoretically explore, the **Levinthal Paradox and Kinetic Partitioning** suggested that proteins sample a subset of fast, directed folding pathways(*1, 2*). The **Energy Landscape Theory** further proposed that proteins explore various conformations while descending a funnel-shaped energy landscape, eventually reaching a thermodynamically stable, low-energy native state(*3*)(*4*)(*5*). Both theories accommodate the diversity of folding pathways but neither clearly define them.

Over time, several models have been proposed to explain the detailed mechanisms of protein folding. The **Framework Model** emphasizes the early formation of local native secondary structures, followed by their diffusion and collision to produce the correct tertiary structure(*6, 7*). In contrast, the **Hydrophobic Collapse Model** proposes a rapid, nonspecific collapse of hydrophobic residues into a disordered core, forming a molten globule, followed by structural rearrangement into the native state(*8*)(*9*)(*10*)(*11*). The **Nucleation-Condensation Model** posits that an extended folding nucleus with weak secondary and tertiary interactions forms first. Around this nucleus, a significant portion of the protein structure condenses into an approximately correct conformation, reaching a transition state before quickly forming the final structure(*12*). Alternatively, the **Nucleation-Propagation Model** suggests that folding starts with the formation of a nucleus consisting of small number of nascent secondary structures. This nucleus then propagates, extending to adjacent sequences until the entire protein adopts its native structure(*13*). Each of these models offers a unique perspective on the folding process, providing insights into how specific proteins achieve their functional conformations.

While some proteins fold spontaneously, many require the assistance of chaperones to reach their native state. The human chaperonin TRiC (also known as CCT), a two-ring stacked complex with eight paralogous subunits in each ring, uses ATP to fold approximately 10% of the proteome(*14*), including the cytoskeletal protein tubulin(*15*). As tubulin emerges from the ribosome in an unfolded state, it is initially captured by the chaperone prefoldin(*16*) and subsequently transferred to TRiC for proper folding(*17*). Upon ATP binding and hydrolysis, TRiC transitions from an open to a closed conformation, encapsulating tubulin within its chamber for folding. In our previous study, we identified three progressive folding intermediates and one final native state of tubulin within the closed TRiC chamber. Electrostatic interactions between the TRiC interior wall and the folded tubulin domains demonstrate that TRiC actively guides tubulin folding as a scaffold, rather than serving merely as a passive Anfinsen cage(*17*). In that study, we defined four non-linear progressively folded domains in tubulin: the N domain (comprising 84% of tubulin’s Rossman fold), the C domain, the core helix domain, and the middle domain (Fig. S1).

Despite this, the tubulin folding nucleus and the mechanism of unfolded tubulin sequence transitioning into the nascent structural elements remain unknown. Here, we combine methods of conventional reconstruction with deep learning to reveal novel tubulin folding states from cryo-EM images of the TRiC/tubulin complex in its closed state, including a subset of particle images previously dismissed as “junk.” We identify the tubulin folding nucleus as a nonnative, partially assembled Rossmann fold. TRiC interacts with the loosely folded structural elements of this nucleus, stabilizing it to facilitate its transition into the first native domain (the N domain mentioned above). We also show that the CCT tails of TRiC promote conformational exploration during tubulin folding, revealing that the folding process is hierarchical, involving nucleation, condensation, and propagation.

## Result

### The compositional heterogeneity in tubulin encapsulation by closed TRiC chamber

The human chaperonin TRiC closes its rings upon ATP binding and hydrolysis to assist with substrate protein folding. However, exactly how the two rings function in a coordinated manner is not well understood. Whether the two rings function independently and are folding active simultaneously remains unknown. To understand how TRiC coordinates its two rings for tubulin encapsulation, we investigate the TRiC binding valency of tubulin.

In our previous study, we induced the ring closure of purified TRiC/tubulin binary complex under ATP/AlFx condition. Focused analysis of the cryoEM closed state particles revealed TRiC rings in the *apo* state (i.e. no tubulin) and rings occupied by tubulin in various folding states, including three intermediate states (I, II, III) and one native state (IV) (Fig. 1A, Fig. S2). Building on that analysis, we trace each ring back to its original 529,181 TRiC particle to determine the compositional heterogeneity within the double-ring TRiC complex (Fig. 1B, 1C). We find a large number of TRiC molecules encapsulate either fully or partially folded tubulin in only one ring, while the opposite ring remains in the *apo* state (Fig. 1C). However, a subset of TRiC molecules encapsulates tubulin in both rings, suggesting that both rings are folding-active and can fold tubulin simultaneously (Fig. 1C). The presence of identical or different tubulin folding states (I through IV) across the two rings illustrates both synchronized and non-synchronized folding in TRiC’s rings (Fig. 1D). This observation suggests the possibility of simultaneous encapsulation, but non-synchronized folding across the rings, or chamber reopening during the folding of the first tubulin to accommodate a second tubulin.

**Fig 1.**
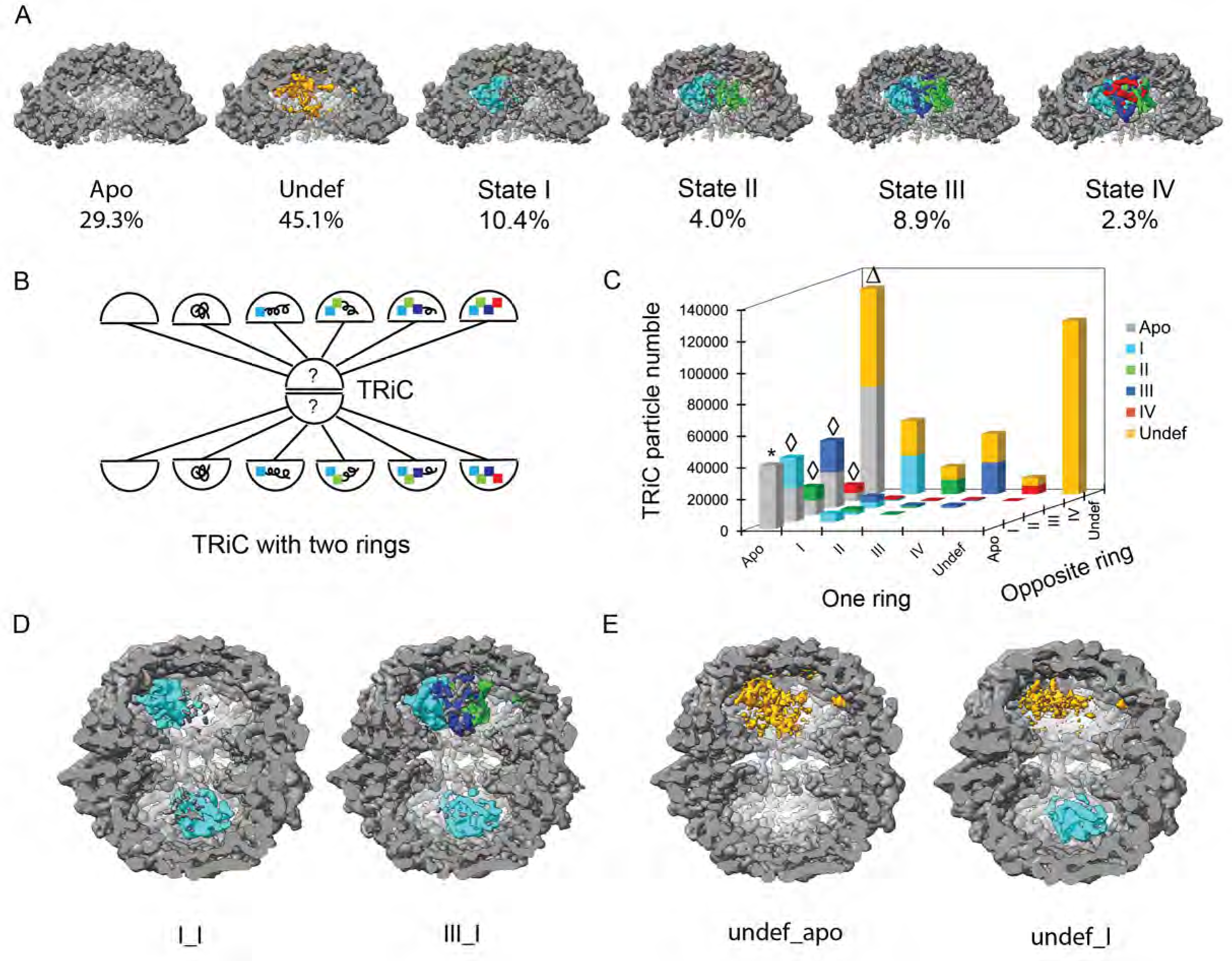
The compositional heterogeneity in tubulin encapsulation by closed TRiC chamber. (A) 529,181 TRiC particles display six types of compositional heterogeneity within their rings, including the apo state, occupied by undefined density, tubulin folding states I, II, III, IV. (B) The diagram shows that each of the two rings in a TRiC can possibly be one of six scenarios in A. (C) The histogram shows the statistical distribution of TRiC particles with compositional heterogeneity from both rings. * The particle set for Fig. 2. Δ The particle set for Fig. 3. ◊ The particle set for Fig. 4. (D) The reconstructions of TRiC with synchronous folding between two rings (left) and the nonsynchronous folding between two rings (right). (E) The reconstructions of TRiC when one ring is occupied by undefined density and the opposite ring is either in the apo state or occupied by a tubulin folding intermediate.

In addition to TRiC rings in the *apo* state or encapsulating one of the four tubulin folding states, a significant portion—45%—of TRiC rings display an undefined density (Fig. 1A,1C). The opposite ring in this population of TRiC can either be in the *apo* state or be occupied by one of the four tubulin folding states (Fig. 1C, 1E). We will characterize this undefined density later in Fig. 3.

### The spatial occupancy and dynamics of CCT tails in apo TRiC

The CCT tails at the inter-ring interface interact with tubulin during its loading and folding process. Here we reconstruct the *apo* state TRiC to 3.7 Å resolution from 40,464 both-rings-*apo* TRiC particle images (Fig. 2A, Fig. S3A). Due to their structural flexibility, many CCT tails are only partially resolvable in the average reconstruction (Fig. S3B). To gain insight into the locations of CCT tails in TRiC, we build the full sequence models of the CCT tails based on the average reconstruction, using two constraints: 1) extend each CCT tail in the direction of its resolved sequence; 2) avoid overlapping with the resolved TRiC density. The density is resolved for the majority of the CCT N tails but less so for C tails (Table S1), thus the structural models for N tails are built with more confidence than those for C tails. From these models, we find that CCT N tails either extend towards the opposite ring (CCTs 1,2,6,7,8) or project out of TRiC through the interring interface (CCTs 4,5) (Fig. 2B, Fig. S3C). In contrast, CCT3’s N tail and all CCT C tails (i.e. CCTs 1,2,3,5,6,7,8; CCT 4 does not have a C tail) likely extend toward the center of the chamber, possibly engaging in intra-ring tails interactions (Fig. 2B).

**Fig 2.**
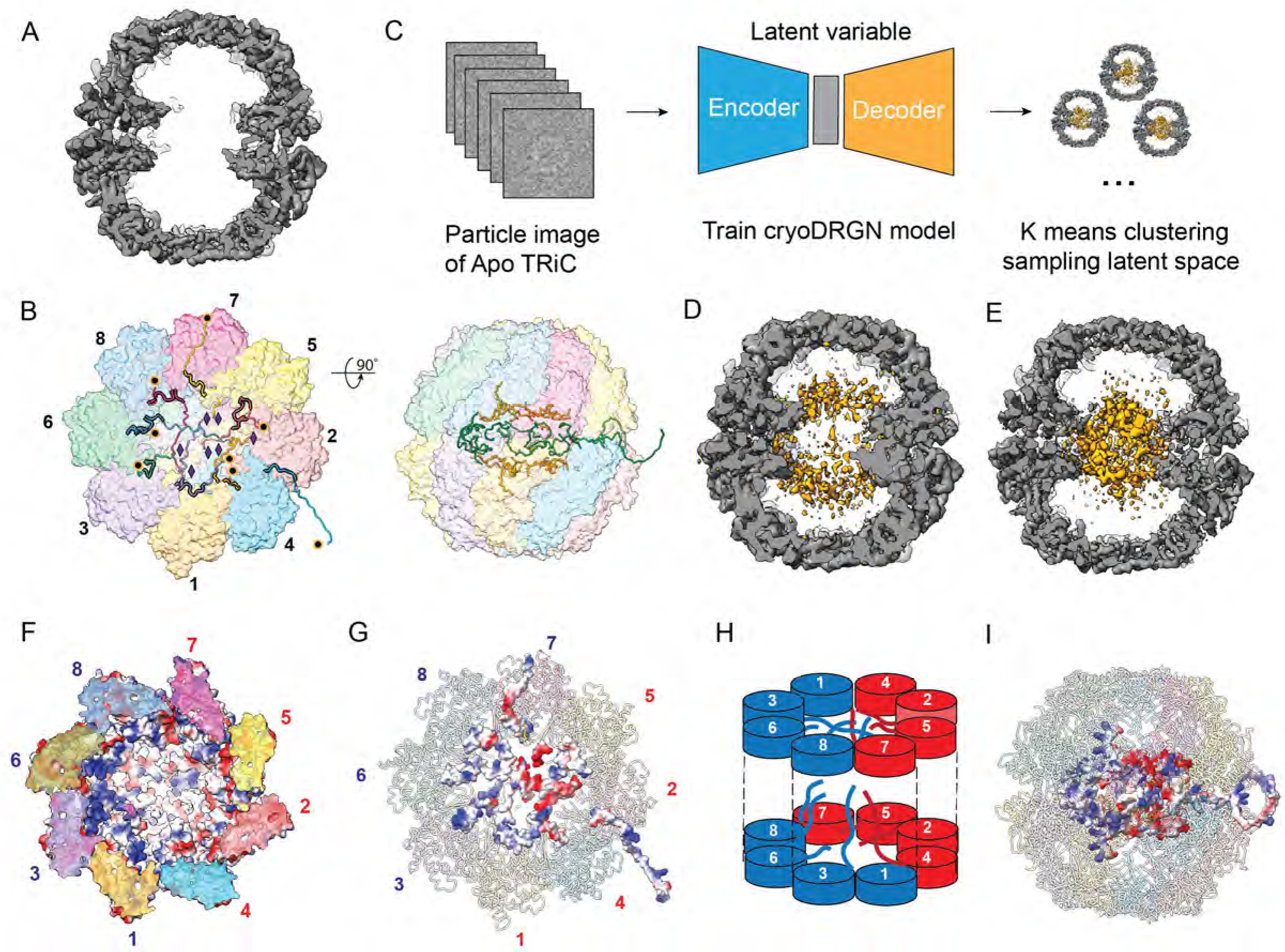
The spatial occupancy and dynamics of CCT tails in apo TRiC. (A) The sliced side view of apo TRiC reconstruction without B factor sharpening. (B) The complete modeling of CCT tails. The left figure shows the CCT tails in a single TRiC ring of the bottom view, with the location of the first N tail residue (orange circle) and the last C tail residue (purple diamond). The resolved CCT tails are highlighted in black stroke. The right figure shows the transparent side view of TRiC with CCT C tails in orange and N tails in green. (C) The workflow of CryoDRGN training and analysis of heterogeneity in apo TRiC particles. (D) The sliced side view of TRiC volume reconstructed from latent space embedding shows the CCT tails density indicating intra-ring termini interaction. (E) The sliced side view of TRiC volume reconstructed from latent space embedding shows the CCT tails density at the inter-ring interface, indicating inter-ring tails interaction. (F) The single TRiC ring model from the bottom view displaying the electrostatic distribution on the TRiC chamber wall. Blue color indicates positively charged distribution while red color indicates negatively charged distribution. (G) The single TRiC ring model from the bottom view displaying the electrostatic distribution on the CCT tails. Blue color indicates positively charged distribution while red color indicates negatively charged distribution. (H) The schematic shows positively charged C tails (CCTs 3,6,8,7) adjacent to positively charged termini (CCTs 1,3,6,8), and negatively charged C tails (CCTs 5,2,4,1) adjacent to negatively charged termini (CCTs 7,5,2,4). (I) The segregation of CCT tails into a positively charged cluster and a negatively charged cluster across the two rings of TRiC.

**Fig 3.**
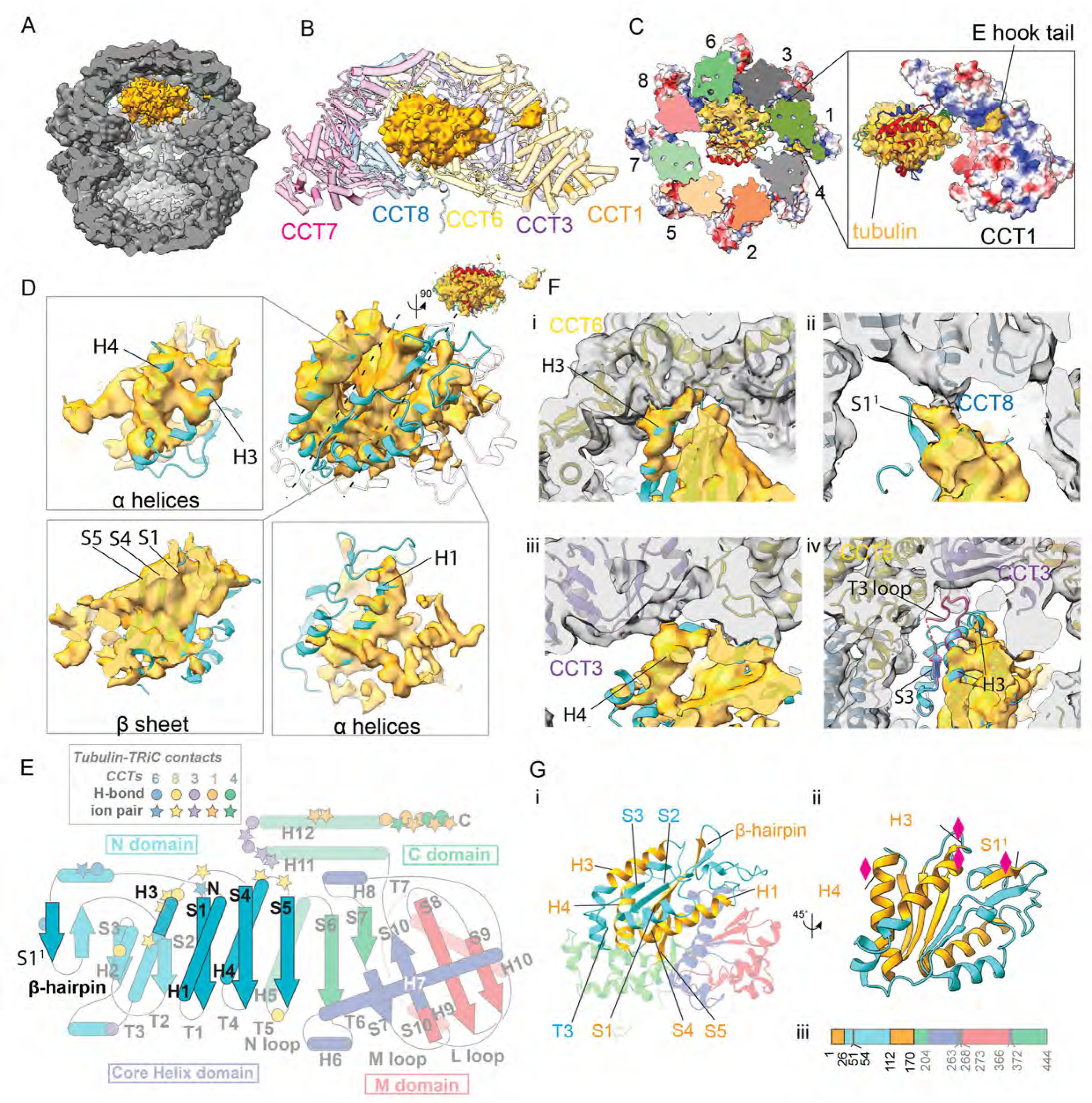
The tubulin folding nucleus State 0.5. (A) The sliced side view of TRiC (in gray color) with one ring encapsulating tubulin density (in orange color) and the opposite ring in the apo state. (B) The tubulin density segmented from the map in A is displayed (in orange color) in the TRiC chamber of CCTs 1,3,6,8,7. (C) The tubulin E hook density at CCTs 1/4 pocket. (D) The three layers of bulky density in State 0.5 (in orange color) correspond to the typical Rossman fold of β sheet sandwiched by helices on its both sides (in cyan color). Figures squared show the density for individual nonnative secondary structural elements in each of the three layers. (E) The topology diagram of tubulin highlighting the loosely folded secondary structural elements (highlighted in cyan) in State 0.5. (F) i, ii, ii show the density of individual nonnative structural elements engaging interaction with CCT subunits and iv shows the absence of T3 loop in State 0.5 (demonstrated by native state model in cyan color), compared to that in State I (model in cornflower blue). (G) i shows the loosely assembled sequence of folding nucleus (in orange) as part of the N domain (in cyan). ii shows the sites of folding nucleus sequence (diamond) engaging interaction with TRiC. Iii shows the folding nucleus (in orange) in one-dimension full sequence of tubulin.

To validate the CCT tail modeling, we extracted the particle stack of *apo* TRiC and investigated the spatial occupancy of CCT tails using cryoDRGN(*18*) (Fig. 2C). K-means clustering and volume reconstruction from the latent space embedding revealed additional density at the inter-ring interface of *apo* TRiC (Fig. 2D, 2E, Movie S1), in contrast to the average reconstruction (Fig. 2A). The extra density observed at the positions of the modeled CCT C tails in many volume reconstructions supports the overall modeling of CCT C tails and their intra-ring contacts (Movie S1, Fig. 2B, 2D). Additionally, density observed across rings at the inter-ring interface in many other TRiC volume reconstructions indicates that inter-ring tails interactions occur in these *apo* TRiC molecules (Fig. 2E, Movie S1). The long, unresolvable C tails of CCTs 1, 3, 7, 8 may cross the ring to participate in these inter-ring tails interactions (Fig. 2H, Table S1).

The closed TRiC chamber comprises a positively charged hemisphere formed by CCTs 1, 3, 6, and 8, and a negatively charged hemisphere formed by CCTs 7, 5, 2, and 4 (Fig. 2F). We observe a similar charge distribution pattern in the CCT C tails as seen on the chamber wall (Fig. 2G). The C tails from neighboring CCTs 3, 6, 8, and 7 form a positively charged cluster near the positive wall hemisphere, while C tails from neighboring CCTs 5, 2, 4, and 1 form a negatively charged cluster near the negative wall hemisphere (Fig. 2F, 2G, 2H, 2I). This electrostatic distribution of CCT tails may play a crucial role in orienting unfolded tubulin by attracting the negatively charged sequence in its N and C domains to the positively charged hemisphere of CCTs 1, 3, 6, and 8 during tubulin recruitment and relocation to the TRiC chamber. In summary, both the modeling and cryoDRGN analysis suggest interactions of C tails both within and between rings in TRiC.

### Tubulin folding nucleus in closed TRiC chamber

The three progressive folding intermediates of tubulin (States I, II, III), along with the fully folded state (IV) (Fig. S1), are situated near or within the native state on the energy landscape(*17*). The polypeptide chain of tubulin, synthesized by the ribosome, collapses into a compact molten globule driven by hydrophobic burial. However, the mechanism by which tubulin folding initiates from the molten globule remains unknown.

To understand the molten globule state of tubulin before a.a. sequence 1-170 (84% tubulin’s Rossman fold) is folded in State I, we examined TRiC particles where one ring was occupied by undefined density while the opposite ring was in the *apo* state, using cryoDRGN. Compositional heterogeneity analysis distinguishes TRiC particles with undefined high density from those with undefined low density (Fig. S4A, S4B). We then reconstruct a 3D map from the pooled particle images with undefined high density in the TRiC chamber (Fig. 3A, Fig. S4C). After fitting the native state tubulin model (State IV, PDB: **7TUB**) to this map (Fig. 3B), we identified the density at the CCTs 1,4 pocket as the tubulin E-hook tail, attributing the undefined density to the full length tubulin (Fig. 3B, 3C).

Using the native state tubulin as a reference, we determined that the N domain of tubulin (i.e. residues 1-170) is partially folded in this reconstruction, hence we designate this tubulin state as State 0.5. In State 0.5, the sequence corresponding to β strands S1/S4/S5 and helices H1/H3(C half)/H4, are loosely folded, representing about half of the six-stranded Rossmann fold (321456) in tubulin (Fig. 3D, 3E, 3G). Their diffuse appearance in the density suggests that these secondary structural elements are in a melted, non-native state.

Although in a non-native state, helix H4, with full density coverage, can be seen to engage with subunit CCT3, while helix H3, partially included in density at its C end, contacts subunit CCT6 (Fig. 3F). While it is unclear whether the assembly of the continuous sequence H3(C half)-S4-H4-S5 occurs before these interactions with TRiC, such interactions could help stabilize the engaged sequences and facilitate the transition from the melted state to fully native structures. The density ascribed to helix H1 is visibly displaced compared to its native state (Fig. 3D). However, the short β-strand S1^1^ of a ® hairpin insertion immediately following helix H1, is included in density at its native position while it interacts with subunit CCT8 as in State I (Fig. 3F). The stabilization of β strand S1^1^ on the TRiC wall could facilitate the formation of the hairpin and likely brings the β strand S2 sequence into proximity with the β strand S1, enabling its annexation. Compared to State I, where the T3 loop is fixed on the TRiC wall(*17*), the absence of the T3 loop density in State 0.5, along with its end motifs–the N half of helix H3 and β strand S3–suggests that the attachment of the T3 loop to the TRiC wall may facilitate the folding of β strand S3. This attachment could help bring the sequence of β strand S3 close to β strand S2, promoting their interaction and annexation (Fig. 3E).

In State 0.5, β strands S1/S4/S5 and helices H1/H3(C half)/H4, although in their non-native states, assemble into a loose tertiary structure that resembles the native state of tubulin (Fig. 3D). This intermediate state supports the nucleation and condensation model of protein folding, with concurrent buildup of non-native secondary and tertiary structural contacts. Consolidation into their native structures, like state I, may require the condensation of additional secondary structural elements in the N domain (Fig. 3G). The nucleation from several secondary structures (H3/H4/S1^1^) contacting the TRiC wall suggests that TRiC actively guides tubulin folding from State 0.5 toward State I (Fig. 3F).

### The propagation dynamics in tubulin folding within TRiC examined in latent space

Understanding the spatial arrangement of the unfolded sequence is crucial for comprehending how a polypeptide transitions from a disordered to a progressively ordered structure and for characterizing the kinetic barriers in the protein folding process. The three tubulin intermediate states (States I, II, III) indicate a domain-wise folding pathway, where discontinuous sequence elements of the tubulin polypeptide fold progressively (Fig. S1). However, the precise mechanism by which sequence of each unfolded domain transitions into its native state during the folding process remains unclear.

To gain insight into the dynamics of the unfolded sequence, we investigate their spatial arrangement in each intermediate state. We pool particle images of TRiC with one ring occupied by each intermediate state while the opposite ring remains in the *apo* state (State x/Apo). We then reconstruct a 3D map of TRiC encapsulating each tubulin intermediate from these pooled particle images (Fig. 4A). Additionally, we embed the pooled images of TRiC encapsulating each state into a separate latent space and reconstruct volumes from the sampled embeddings to examine the conformational heterogeneity in the unfolded sequence for each intermediate state (Movies S2, S3, S4, S5, Methods).

**Fig 4.**
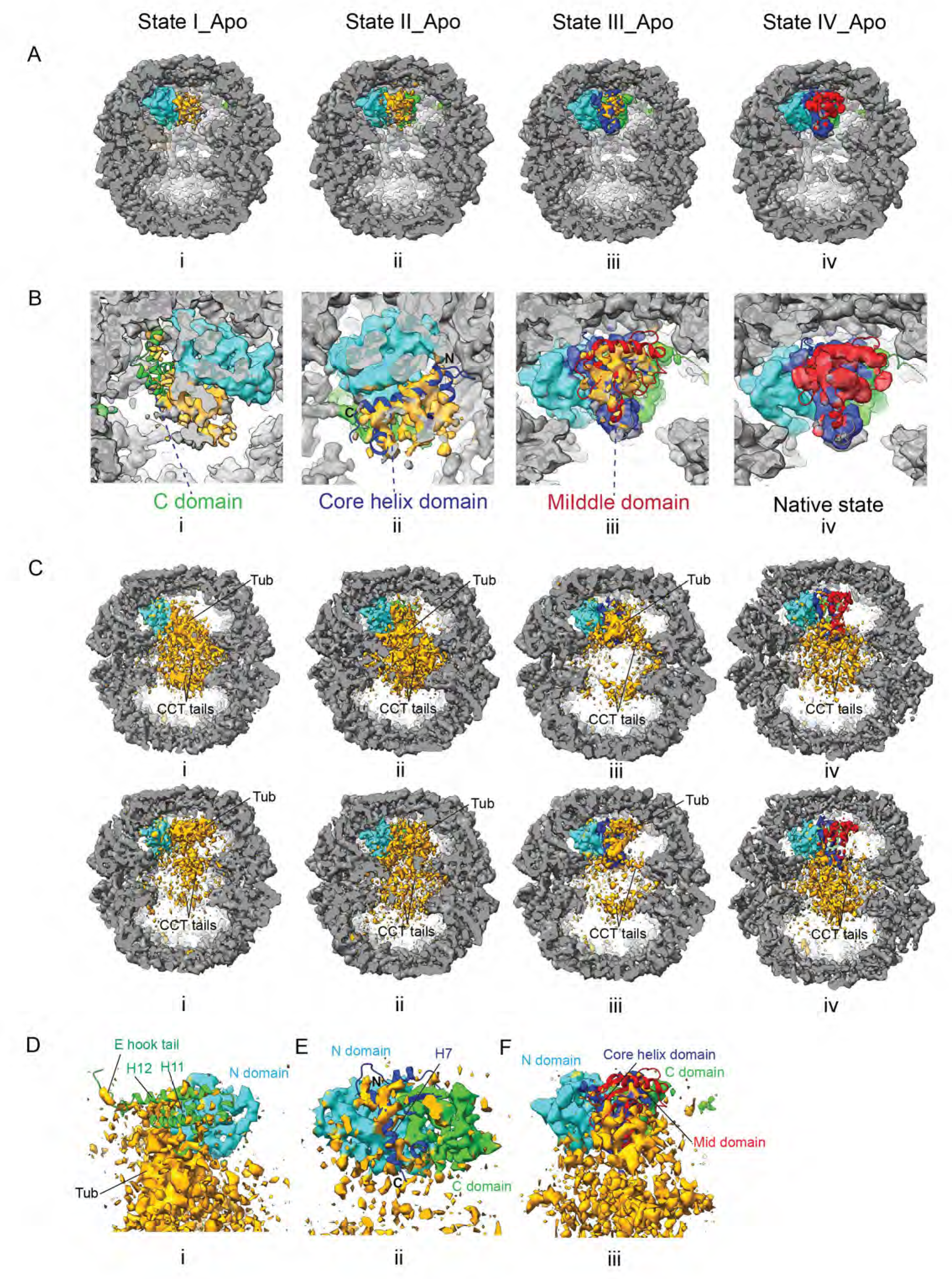
The propagation dynamics in tubulin folding within TRiC. (A) The reconstructions of TRiC with one ring encapsulating tubulin intermediate State I, II, III or native state IV, while the opposite ring is in the apo state. (B) Zoom in view of tubulin density in (A). i shows the C domain is barely covered by density in State I; ii shows the helix H7 in Core helix domain is covered by visibly displaced density in State II; iii shows the Middle domain density in state III condensates around native position; iv shows the fully folded tubulin density in State IV covers the secondary structures of native state tubulin. (C) The examples show the density for tubulin unfolded sequences and the CCT tails in the TRiC chamber from state I to State IV by cryoDRGN analysis. (D) The example cryoDRGN volume indicates the density is displaced from the C domain helices H11 and H12 in State I; (E) The example of cryoDRGN volume shows the helix H7 from core helix domain is partially covered by density in State II; (F) The example of a cryoDRGN volume indicates the density of unfolded middle domain is displaced from the native position in State III.

In the average reconstruction of State I/Apo, the folded N domain, accounting for 38% of the tubulin sequence, is well-resolved (Fig. 4A, Fig. S1). However, the average density for the remaining 62% unfolded sequence appears weak due to heterogeneity (Fig. 4B). Superimposing the native state tubulin model shows that the C domain, which appears folded in State II, is barely covered by density (Fig. 4B), indicating that sequence of this domain infrequently occupies the native state position among the analyzed particles in State I/Apo. Latent space sampling reveals that the unfolded sequence undergoes extensive spatial rearrangement, despite being constrained by the TRiC chamber (Movie S2, Fig. 4C). This diverse spatial arrangement enhances the conformational space sampling of the unfolded sequence. The flexible CCT tails from the same or across chamber perturb the unfolded sequence through direct contacts (Movie S2, Fig. 4C), likely driving it out of local energy minima. Examination of the latent space volumes for State I/Apo reveals the absence of resident density for the C domain helices H11 and H12 at their native positions, although the tubulin C-terminal E-hook tail immediately following these helices is observed still attached to the CCT1/4 pocket of TRiC (Movie S6, Fig. 4D). Condensation of C domain to the approximate native position seems challenging and given the relatively small percentage of State II among the folding intermediates (Fig. 1A), the final folding of the C domain to State II is likely a rate-limiting step in the folding process.

In the average reconstruction of State II/Apo, further folding of the C domain leaves 37% of the tubulin sequence still unfolded (Fig. 4A, Fig. S1). However, diffuse density is observed at the position of native helix H7 in the tubulin Core helix domain, which appears folded in State III, although it is visibly displaced toward its C end (Fig. 4B). Latent space volumes of State II/Apo reveal that the unfolded sequence remains highly dynamic within the TRiC chamber (Movie S3, Fig. 4C). Despite the overall extensive movement of the unfolded sequence, persistent density coverage is observed at the N end of helix H7, indicating a non-native fold in this region (Movie S7, Fig. 4E). In contrast, the C end of helix H7 shows variable density coverage among the volumes (Movie S7), indicating the C end is actively condensing on the surface of the folded N/C domain. The folding of helix H7 propagates from the N to the C end, arguing against the framework model which posits pre-established secondary structures. Instead, the folding of the Core helix domain is consistent with the canonical nucleation and propagation model of protein folding, involving the gradual formation and growth of secondary structures around a native nucleus. Compared to the C domain in State I/Apo, helix H7—the major secondary structure of the Core helix domain—restricts the spatial search to the vicinity of its native state position, suggesting an easier condensation of Core helix domain, likely leading to a fast progression from State II to State III, supported by the relatively high percentage of State III among the folding intermediates.

In the average reconstruction of State III/Apo, where the Core helix domain is further folded, density corresponding to the remaining 14% unfolded sequence is observed around the position of the Middle domain of native state tubulin (Fig. 4A, 4B, Fig. S1). Many latent space volumes of State III/Apo also reveal similar spatial arrangement in the unfolded density (Fig. 4C, Movie S4). However, many other volumes have the unfolded density located away from the native state position (Fig. 4C, 4F, Movie S4). The flexible CCT tails may help propel the unfolded sequence toward the native state position for folding. Overall, the unfolded sequence has a more restricted search space due to its shorter length and stabilization at both ends by the already folded regions (Fig. S1).

In the average reconstruction of State IV/Apo, the final folding of the Middle domain clears the TRiC chamber (Fig. 4A, 4B). The latent space volumes for State IV/Apo align with the average reconstruction, though occasional residual density likely corresponding to the flexible CCT tails is observed elsewhere in the chamber (Fig. 4C, Movie S5). In these latent space volumes, the CCT tails are observed below the tubulin at the inter-ring interface, engaging the intra-ring or inter-ring tails interaction (Fig. 4C). Throughout the progressive folding of tubulin from State I until its native State IV, the CCT tails consistently interact with the tubulin sequence, facilitating the conformational sampling by spatial rearrangement (Fig. 4C).

## Discussion

Folding nuclei (FN) refers to a minimal compact region of protein at the early stage of the folding process that establishes a framework for subsequent rapid assembly of the native state. The existence and nature of FN are central to understanding how proteins fold. Over the past few decades, experimental identification of FN has been attempted using several techniques, including Φ-value analysis(*19*)(*12*)(*20*), Nuclear Magnetic Resonance (NMR) spectroscopy(*21*), Single-Molecule Fluorescence Resonance Energy Transfer (smFRET)(*22*) and Single-molecule force spectroscopy(*23*). Though they are indirect methods with limitations such as the unknown impact of mutations, resolution, size limitations, complexity of multi-domain proteins, and signal overlap, these techniques provide important insights about the FN. In contrast, theoretical experiments allow simulation of the protein folding process to directly visualize the FN by molecular dynamics (MD) Simulations, Coarse-Grained Simulations and Energy Landscape Modeling. However, issues such as time scale limitations, force field accuracy, conformational sampling, solvent effects, and computational cost make the accurate prediction of the location of the FN challenging for even small proteins.

The tubulin polypeptide collapses into a molten globule state during ribosome translation. However, the mechanisms of folding initiation and propagation remain unclear. In this study, we directly visualize the folding nucleus as a loosely and partially assembled Rossman fold by cryo-EM. The assembly of non-native secondary structural elements into a tertiary structure resembling part of the native state Rossman fold suggests the folding nucleus as a dry molten globule state(*24*)(*25*). The canonical nucleation-condensation model posits that a metastable and short-lived nucleus attracts rapid condensation of a large portion of the sequence around it, leading to the native state. However, our observation of the tubulin folding nucleus within a large population of TRiC molecules contrasts this assumption. TRiC contributes to stabilizing the folding nucleus by interacting with its non-native structural elements. The condensation of the remaining N domain sequence into an approximately correct conformation is expected to transition the full N domain into its native state, as observed in State I.

The Rossmann fold is one of the most primitive protein folds, emerging early in evolution and found across all domains of life, including bacteria, archaea, and eukaryotes. Its simplicity and stability likely facilitate spontaneous and efficient folding, as exemplified by proteins like the tubulin paralog FtsZ in E. coli. However, in tubulin, the Rossmann fold does not fold spontaneously during ribosomal translation, as indicated by intermediate State 0.5. An additional sequence insertion consisting of a β hairpin between helix H1 and β strand S2 may hinder the rapid condensation of β strand S2 onto the folding nucleus, thereby possibly impeding the proper folding of the N domain. Notably, this insertion is partially stabilized in intermediate State 0.5 and further stabilized in State I by the TRiC chaperonin complex. This suggests that TRiC assists tubulin in achieving the native folding of the N domain by sequestering the insertion sequence away from the key structural elements of the Rossmann fold.

Both the average reconstruction and volume reconstructions generated from latent space embedding reveal the complete relocation of the tubulin sequence from the inter-ring space to the TRiC chamber upon TRiC ring closure. In State 0.5, the TRiC chamber is almost entirely occupied by unfolded tubulin, which would complicate a large reorientation if the N and C domain sequence were initially positioned incorrectly, e.g. facing hemisphere of CCTs 7/5/2/4 instead. This observation supports our hypothesis that the electrostatic distribution in the CCT tails plays a crucial role in orienting the unfolded tubulin by attracting the negatively charged sequences of the N and C domains to the CCT1/3/6/8 hemisphere during the tubulin recruitment and relocation stage.

We observe double occupancy of tubulin within the closed TRiC, indicating simultaneous folding in both rings. This contrasts with previous studies in which plp2 occupies one ring to assist tubulin folding in the opposite ring(*26*). Three hypotheses that could explain this observation warrant further investigation: 1) plp2 may be dispensable but significantly accelerates tubulin folding; 2) plp2, as observed in previous studies, may be on standby to assist in the folding of actin rather than tubulin; and 3) plp2 may participate in tubulin folding specifically when TRiC is in its open state.

Protein folding, evolution, and de novo design are interconnected fields that rely on a thorough understanding of how a polypeptide’s primary structure dictates an efficient folding pathway, ultimately leading to a well-defined tertiary structure. Protein folding—whether spontaneous or chaperone-mediated—remains a frontier in molecular biology. In this study, we move beyond static snapshots of tubulin folding intermediates to describe the condensation and propagation of individual structural elements within the folding nucleus (State 0.5) and on the natively folded domains (States I, II, III), providing detailed insights into the tubulin folding pathway. Additionally, we examine the range of spatial arrangements of the unfolded sequence in each intermediate state within the TRiC chamber, revealing that CCT tails actively promote the conformational space sampling during tubulin folding. These observations lay the groundwork for future computational experiments on tubulin folding and drug design.

## Author contributions

Conceptualization: Y.Z., W.C.; Methodology: Y.Z.; Investigation: Y.Z.; Writing – original draft: Y.Z.; Writing – review & editing: Y.Z., M.S., W.C.; Funding acquisition: W.C..

## Data and materials availability

Maps have been deposited in the Electron Microscopy Data Bank (EMDB) with the accession codes EMDB XXXX.

## Supplementary Materials Materials and Methods Supplementary Text

Figs. S1 to S4

Tables S1

Movies S1 to S7

